# Targeted Molecular Dynamics Simulations Suggest Direct Ligand Competition as a Plausible Efflux Inhibition Mechanism in the Multidrug Resistance Pump AcrB

**DOI:** 10.1101/339242

**Authors:** Lande Silva, Pedro Eduardo Almeida da Silva, Karina S. Machado, Nelson Dutra, Terry P. Lybrand

## Abstract

We report computer simulation results using the Targeted Molecular Dynamics technique to explore possible transport mechanisms in the multidrug efflux pump AcrB for two substrates, ethidium bromide and a tetrahydropyridine derivative. These studies revealed structural elements, including specific α-helices, β-strands and flexible loops that define a physically plausible pathway for substrates to the extracellular environment. These calculation results can be used to plan future biophysical experiments and may suggest interesting drug design possibilities to address drug resistance due to AcrB function.

**Importance:** Addressing the issue of antimicrobial resistance mediated by efflux, this study presents possible binding sites and structures in the AcrB MDR pump that could be molecular targets for drugs. Targeted molecular dynamics simulations suggested that these sites and structures seem vital for a successful efflux. The AcrB is proposed to be divided into three distinct zones, with loops, sheets and helices mediating the passage of molecules from one zone to another. We also described possible capture sites on the outer part of the protein and access ways to its interior. Finally, we proposed that ligand competition for same pathways could be thought as an efflux inhibitory mechanism, thus assisting to conceive new ways of designing efflux pump inhibitors.

## 1. Introduction

The multidrug efflux pump AcrB from *Escherichia coli*, a member of the resistance-nodulation division (RND) family transporters, has been studied extensively as a model for RND efflux pumps that occur in gram-negative bacteria. It is responsible for the capture and extrusion of a wide variety of substrates, like dyes, heavy metals and antibiotics from the cell (1). These efflux pumps contribute to bacterial resistance for many antimicrobial agents and biocides (2), and thus constitute a major public health concern (3). Therefore, it is important to obtain a more detailed understanding of the AcrB ligand capture-extrusion mechanism, as this information may suggest effective strategies to inhibit the efflux process, thus improving the efficacy of antimicrobials and reducing drug resistance in many gram-negative microorganisms.

Targeted Molecular Dynamics (TMD) is a method that induces conformational changes in a structure based solely on constraints applied to minimize the root mean square deviation between initial and final (target) structures (4). TMD has been used previously to examine protein conformational changes induced by ligand binding (5) and to explore ligand binding reaction coordinates (6). The only information necessary to perform TMD calculations are detailed three-dimensional structures for the complex in both an initial (I) and a final (F), or target, state. The I and F states are usually obtained from x-ray diffraction or NMR studies for the ligand-protein complex. If structures are available only for the unliganded protein, as is the case for AcrB in this work, plausible I and F states can be generated using molecular docking calculations. TMD calculations were performed for both ethidium bromide (EtBr) and a tetrahydropyridine derivative NUNL02 (7)(8) (Fig. A1) to characterize possible transport pathways in AcrB for the ligands from the intracellular surface to the TolC domain.

**Fig. 1.**
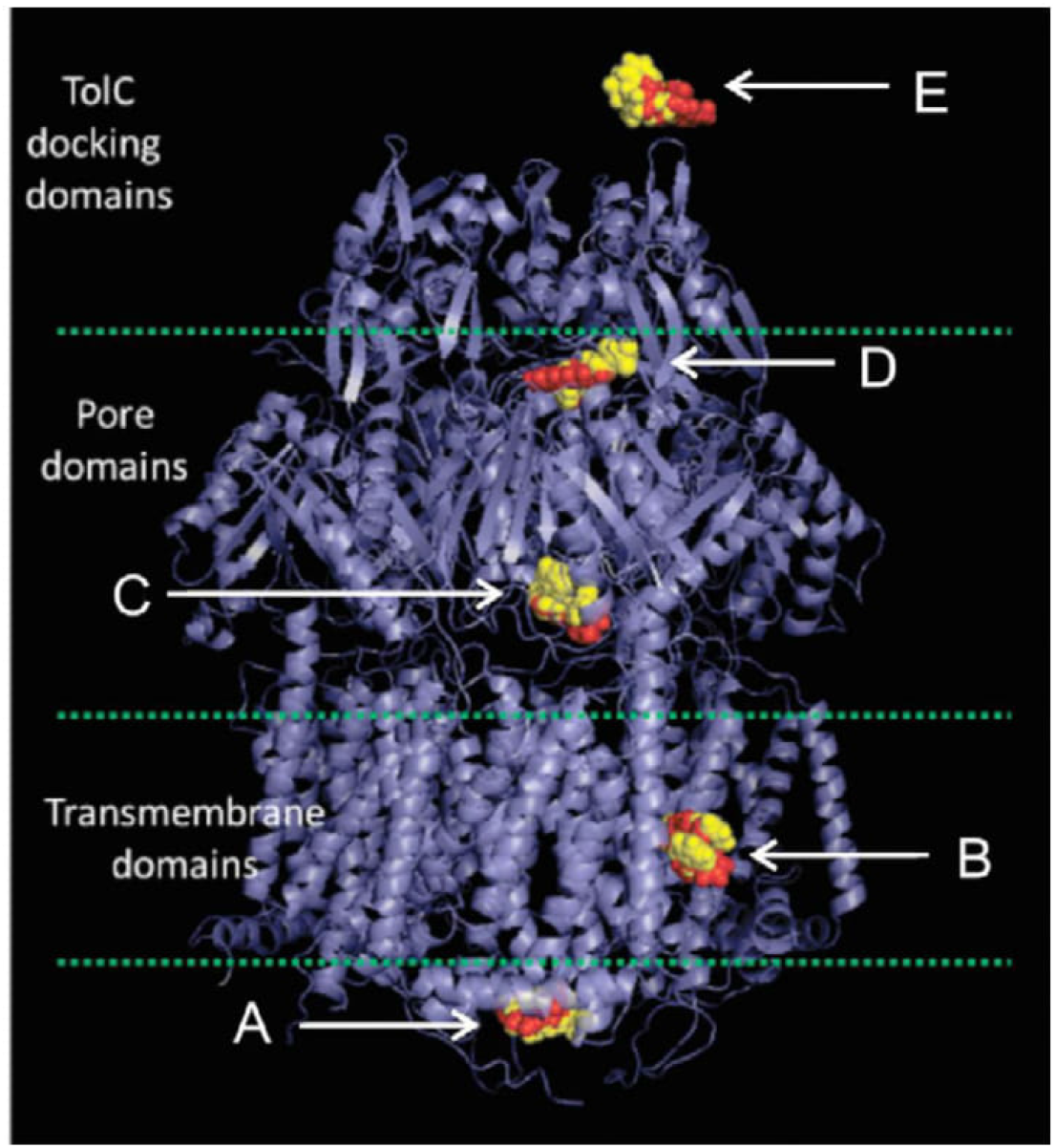
An overview of the positions chosen for the TMD procedure. EtBr is depicted as red CPK models and NUNL02 as yellow CPK models while the AcrB protein is displayed as a blue ribbon structure. Position A is located at the cytosolic surface of the transmembrane domain, position B is at the transmembrane domain-lipid membrane interface, position C is at the edge of the pore domain, position D is at the boundary between the pore and the TolC domains, and position E is outside the TolC domain in the periplasmic space.

The TMD simulations revealed interesting conformational changes in the protein backbone as the ligands progressed through AcrB from the cytosolic interface to the periplasmic surface. These results reinforce the idea of competition as a mechanism of efflux inhibition discussed in (9), in the present work, however, different ligands seem to interfere with transport of each other by utilizing the same transit “pathways” through the protein structure.

## 2. Material and Methods

### 2.1 Docking

The AcrB structure without bound ligands (PDB ID: 1IWG) (10) is considered to be in the resting state for the transporter and was chosen for the TMD studies. This ligand-free, symmetric homotrimer structure has a resolution of 3.5 Å, and 1053 residues in each subunit. The structure is divided into three domains: a transmembrane domain, a pore domain and the TolC interaction domain that together constitute the periplasmic headpiece (10) (11) (Fig. 1). Some short loop segments, between residues 496 and 513, 708 and 716 and 858 and 871, were not resolved in the x-ray structure, and we used Modeller 9.14 (12) (13), with default parameters, to construct them.

Experimental results suggest that both EtBr and NUNL02 are substrates for the multidrug efflux pump AcrB (9) (11) (14). Following the docking methodology described in (9), we used Autodock Tools (15) and AutoDock Vina (16) to locate high-affinity binding sites for the substrates EtBr and NUNL02 in chain A of the full AcrB model, ranked by free energy of binding (FEB) scores. Dockings were performed with the exhaustiveness set to 8 (run 1), and later set to 128 (search grid sizes of 40 × 48 × 7 and 65 × 65 × 23, respectively) (run 2). The grids were moved from immediately below the transmembrane domain to the TolC domain, in steps of 5 Å in both cases, as previously described (9). Positions A, B, C, D and E (Fig. 1) were determined from run 1 and confirmed with run 2 and by comparison with dockings performed for structure PDB ID: 4DX5-A (17). No significant differences for determining the I and F positions for the TMD were found between the docking runs (Fig. A2 to A5). The docking calculations generated five favorable binding sites in the AcrB structure, including positions A and B in the transmembrane domains, positions C and D in the pore domain and position E in the TolC domain (Fig. 1) (Table 1). Position A had reasonable FEB scores in run 1 and 2 at the AcrB surface at the cytosol-membrane interface, and we selected this location to represent the initial state for modeling ligand capture and efflux directly from the cytosol to the periplasmic region. Position E does not have a particularly favorable docking score compared to the other selected docking poses, but was chosen to represent a position completely outside the AcrB structure, i.e., a position for an extruded ligand, and we used the position E complex as the F state. Since AcrB forms a complex with the TolC protein (11), the ligand would likely be bound to the TolC protein at position E. Since the 1IWG crystal structure presumably represents the inactive state of the transporter, it is possible that these predicted ligand binding sites might not be mechanistically relevant. However, previous molecular docking studies (9) using the AcrB 4DX5 chain A crystal structure (17), presumed to represent an active state of the transporter, yielded EtBr and NUNL02 binding sites that correspond closely to the binding sites identified in the current study (Fig. A6 to A9). Therefore, we used the five positions illustrated in Fig. 1, as initial and final states for a series of TMD simulations to explore plausible ligand efflux pathways. A symmetric homotrimeric structure was then generated using the full chain A model with Pymol (18). Figures were made with VMD (19) and Pymol (18).

**Table 1.** Free energy of binding (FEB) found in run 1, for the positions that compose the pathways for EtBr and NUNL02.

|  | Position | A | B | C | D | E |
| --- | --- | --- | --- | --- | --- | --- |
| Energy<br>(kcal/mol) | EtBr | -5.8 | -7.1 | -8.2 | -7.4 | -3.2 |
|  | NUNL02 | -7.2 | -6.2 | -7.6 | -6.8 | -1.1 |

### 2.2 Targeted Molecular Dynamics

We chose the TMD method for this study because this technique does not require an explicit definition of a detailed reaction pathway or restraint coordinates that could potentially bias results if inappropriate restraints were applied. The full efflux pathway was sub-divided into discrete steps, each probed with individual TMD simulations, to facilitate study of alternate possible pathways.

Pathway 1: In this pathway, either EtBr or NUNL02 was placed initially at position A. Sequential TMD simulations were then performed to follow ligand transit from position A to position C, then position C to position D, and finally position D to position E outside the AcrB protein (Fig. 1 and 2). We did not attempt to model detailed mechanisms for the initial binding of either ligand to position A. Position A simply represents a plausible site for initial ligand binding if AcrB captures the ligand directly from the cytoplasm as suggested in a previous study (10).

**Fig. 2.**
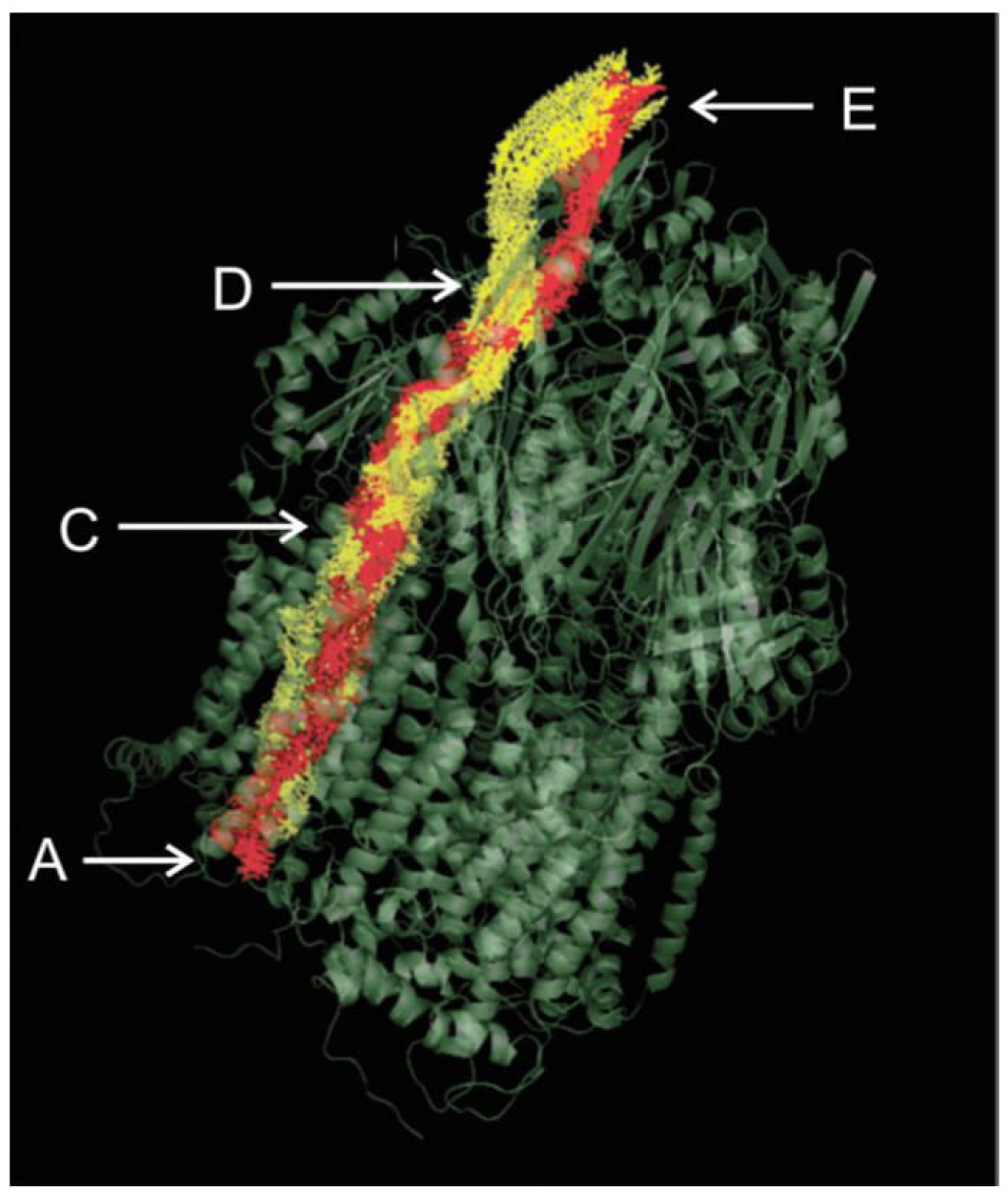
The EtBr (red pathway) and NUNL02 (yellow pathway) follow the same path inside the AcrB efflux pump, from position A to C to D. However, the ligand paths begin to diverge after position D.

Pathway 2: In this pathway, either EtBr or NUNL02 was placed initially at position B. Position B represents a plausible initial ligand binding site if AcrB captures the ligand from the membrane domain rather than the cytoplasm as proposed by Nikaido, et al. (20). TMD simulations were then run to follow ligand migration from position B to position C. After this point, all Pathway 2 details are identical to Pathway 1.

All simulations were performed using the AMBER 12 package (21). Atomic charges and any additional missing parameters for EtBr and NUNL02 were generated using the ANTECHAMBER utility in AmberTools14 (22). The xLeap module was used to add missing hydrogens and Na+ counterions to neutralize the full complex, and the system was solvated in a truncated octahedron water box. Since the periplasmic headpiece (the pore domain and the TolC interaction domain) constitute the majority of the AcrB transporter, we decided to perform solution phase simulations rather than construct a more complicated aqueous bilayer model to embed the transmembrane domain. We monitored the transmembrane domain helical bundle during the simulations to confirm its structural stability during the solution phase simulations. To ensure that the initial and final states for each TMD simulation contained identical numbers of water molecules, the solvation calculations were performed with ligand present at both the initial (position A or B) and final (position E) sites. A single ligand was then deleted as appropriate to generate either the initial or final state model, respectively.

For each solvated complex, the protein and ligand atom positions were constrained while water molecules and counterions were relaxed with 500 steps of steepest decent minimization followed by 500 steps of conjugate gradient minimization using an 8 Å nonbonded cutoff. Next, the full protein-ligand complex along with solvent and counterions was relaxed with 1000 steps of steepest descent and 1500 steps of conjugate gradient minimization using a 10 Å nonbonded cutoff and particle-mesh Ewald corrections for long-range electrostatics (23). The ff12SB and gaff force fields were used, respectively, for protein and ligands. Then, each solvated complex was heated slowly from 0 to 300K during a 20 ps NPT ensemble MD simulation, with protein heavy atoms weakly restrained at the minimized structure. The 300K NPT MD simulation was propagated for an additional 100 ps with no positional results to generate starting configurations for each TMD calculation. All TMD simulations were performed using both 0.5 kcal/mol-Å and 1.0 kcal/mol-Å force constants to assess possible biasing effects of restraint force constant choice.

## 3. Results

### 3.1 Molecular Docking

Molecular docking calculations were performed independently for each ligand. Fig. 1 highlights the significant overlap for EtBr (red CPK) and NUNL02 (yellow CPK) at all five binding sites, suggesting the possibility of direct efflux competition between these substrates, as these binding sites represent stable intermediate states along the simulated efflux pathways. As noted above, molecular docking calculations using either the 1IWG crystal structure (inactive conformation) or the 4DX5-A crystal structure (active conformation) yielded very similar binding poses for all five positions depicted in Fig. 1, so the docking results do not appear to be particularly sensitive to the exact protein conformation, as least for these five binding sites.

### 3.2 Efflux Pathways

#### 3.2.1 Pathway 1

The TMD results for this pathway show that both ligands follow essentially the same path as they traverse from position A to position D through the AcrB protein, as displayed in Fig. 2 and in the animated movies A.10 and A.11. Interestingly, from position D to position E, the EtBr and NUNL02 paths begin to diverge significantly. Detailed analysis of pathway 1 for EtBr efflux suggests a gated transit mechanism through an extended tunnel with constriction points that open transiently, apparently as result of specific interactions with the ligand. Dividing pathway 1 into three distinct zones made it easier to identify residues and structural elements that appear to play an important role in the substrate efflux mechanism (Fig. 3).

**Fig. 3.**
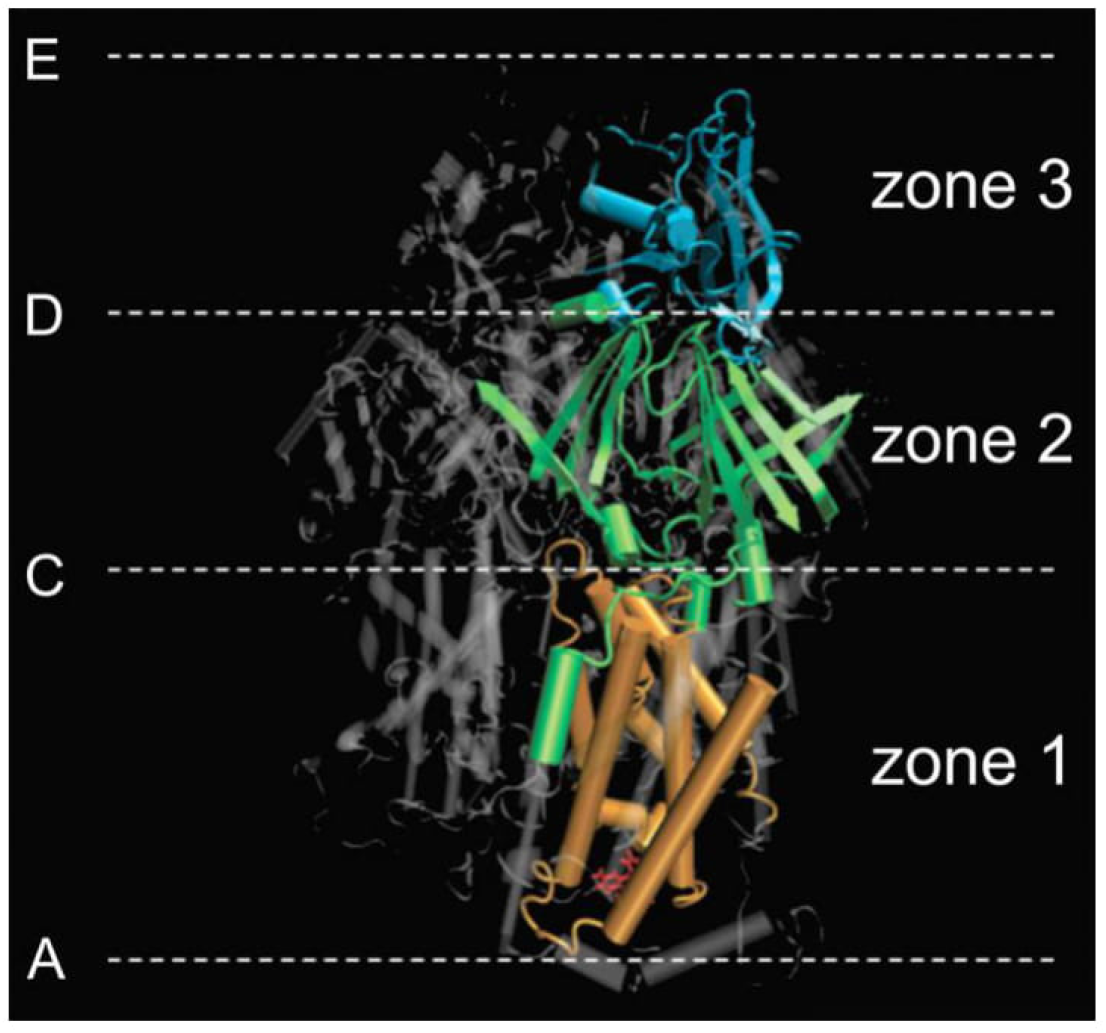
Pathway 1 zones. Orange: Zone 1, between position A and position C, comprises much of the transmembrane domain. Green: Zone 2, between position C and position D, consists primarily of interior β-strand structure in the pore domain. Blue: Zone 3, between position D and position E, consists of the TolC interaction domain.

Zone 1 (Fig. 4) is localized to the transmembrane domain. This region has a total of 12 transmembrane α-helices (10), 9 of which form an apparent transit tunnel through the transmembrane domain from the cytosolic surface to the boundary with the pore domain. These nine helices are displayed in Fig. 4. Table 2 lists the residues in each helix.

**Fig. 4.**
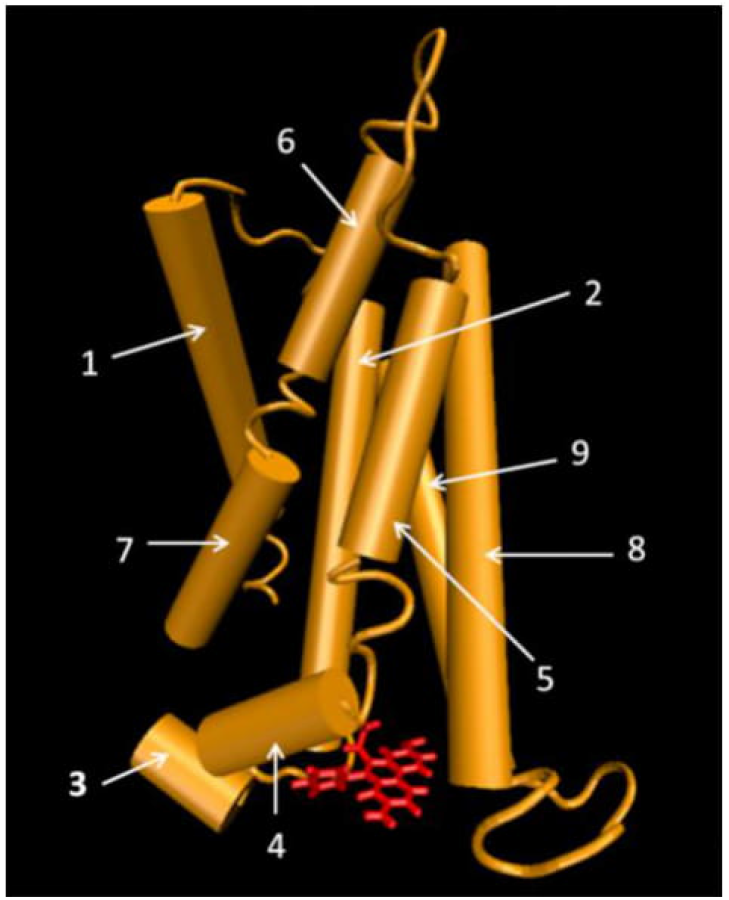
Zone 1 from pathway 1. The nine α-helices that form a tunnel structure in the transmembrane domain are displayed in orange. EtBr (red) is displayed in the position A binding site at the protein cytosolic surface.

**Table 2.**
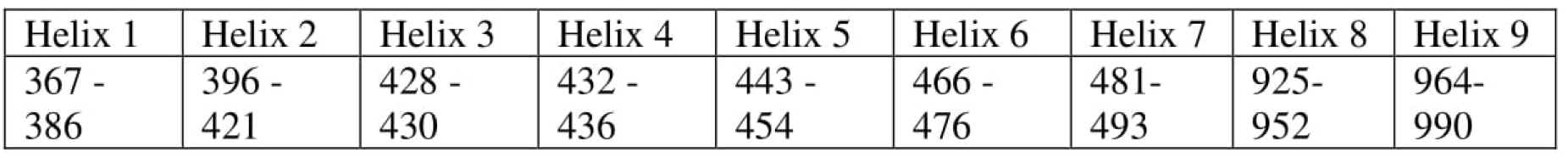
Zone 1 helix residues displayed in Fig. 4.

Position A (Fig. 5a) identified in the molecular docking calculations is the initial ligand binding site for pathway 1. Initial ligand binding at this position would enable the AcrB efflux pump to transport molecules directly from the cytoplasm. Ligand transport through Zone 1 is correlated with a peristaltic motion of the nine α-helices that form the transient tunnel, as displayed in Fig. 5. Initially, the helices are packed tightly when the ligand binds at position A. As the ligand enters Zone 1, the helical bundle relaxes (Figs. 5b and 5c), providing a transient passageway for the ligand to navigate through the transmembrane domain. As the ligand exits Zone 1 to occupy position C (Fig. 5d), the helical bundle reverts to the tightly packed conformation observed before ligand entry.

**Fig. 5.**
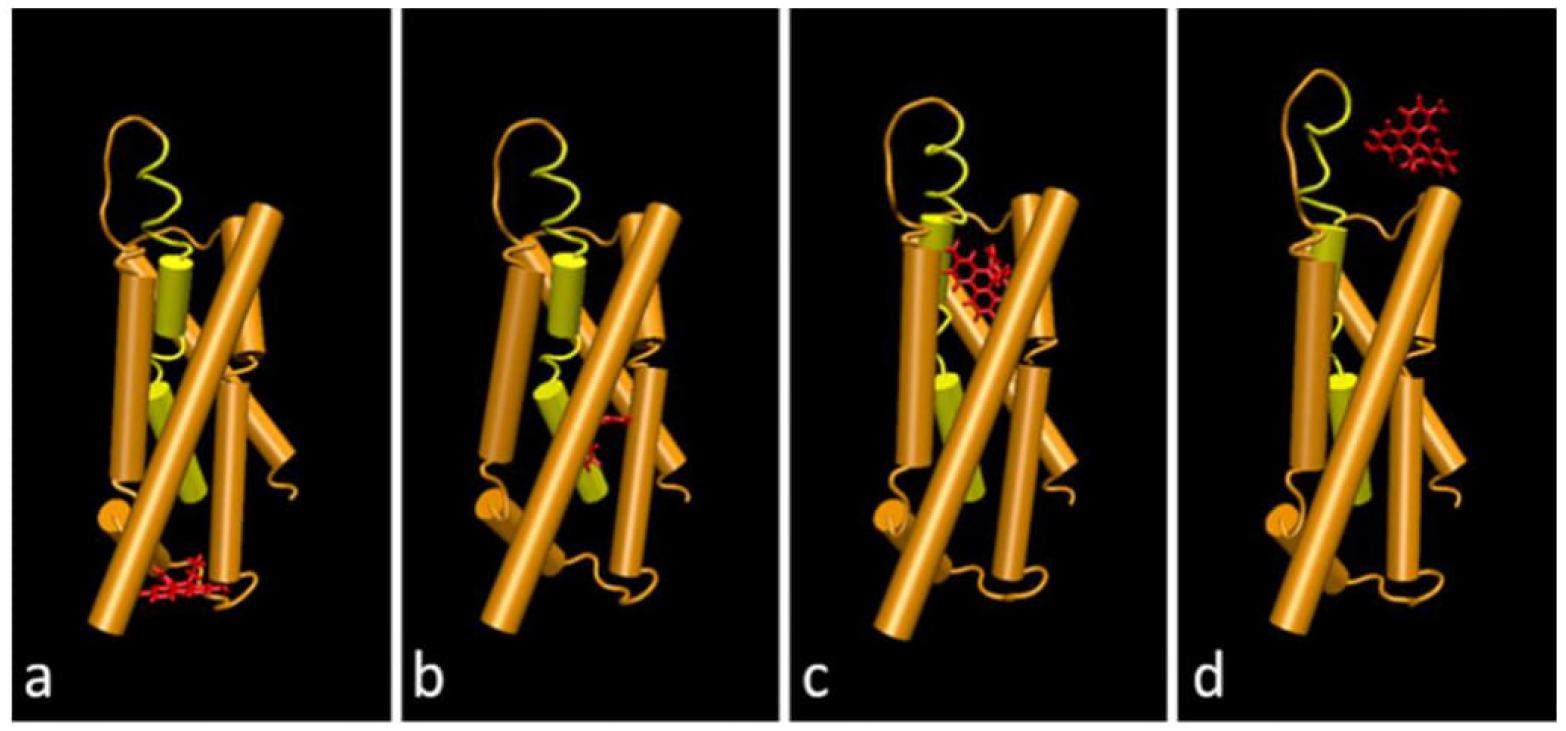
Helix bundle movements in Zone 1 that are coupled to ligand transit. Helices 6 and 7 are displayed in yellow. Panel a: EtBr is bound at position A and the helix bundle is tightly packed; Panels b and c: helical shifts reduce the tight bundle packing, allowing EtBr to enter the transiently opened passageway; Panel d: as EtBr exits Zone 1, the helical bundle resumes its original tight-packing arrangement.

It appears that these conformational changes are induced by ligand interactions, as these conformational fluctuations are not observed in the Zone 1 helical bundle in the absence of ligand. The two helices displayed in yellow in Fig. 5 correspond to helices 6 and 7 (Fig. 4). In fact, these helices may be a single helix with a flexible “elbow” segment at residues 478-480. Backbone conformational changes at these residues, induced by the ligand, result in a significant reorientation of helical segment 6 relative to helix 7. The reorientation of helix 6, along with more modest shifts for helices 1 and 2 (Fig. 6), reduce the tight helix bundle packing and generate a transient passageway large enough for the ligand to navigate.

**Fig. 6.**
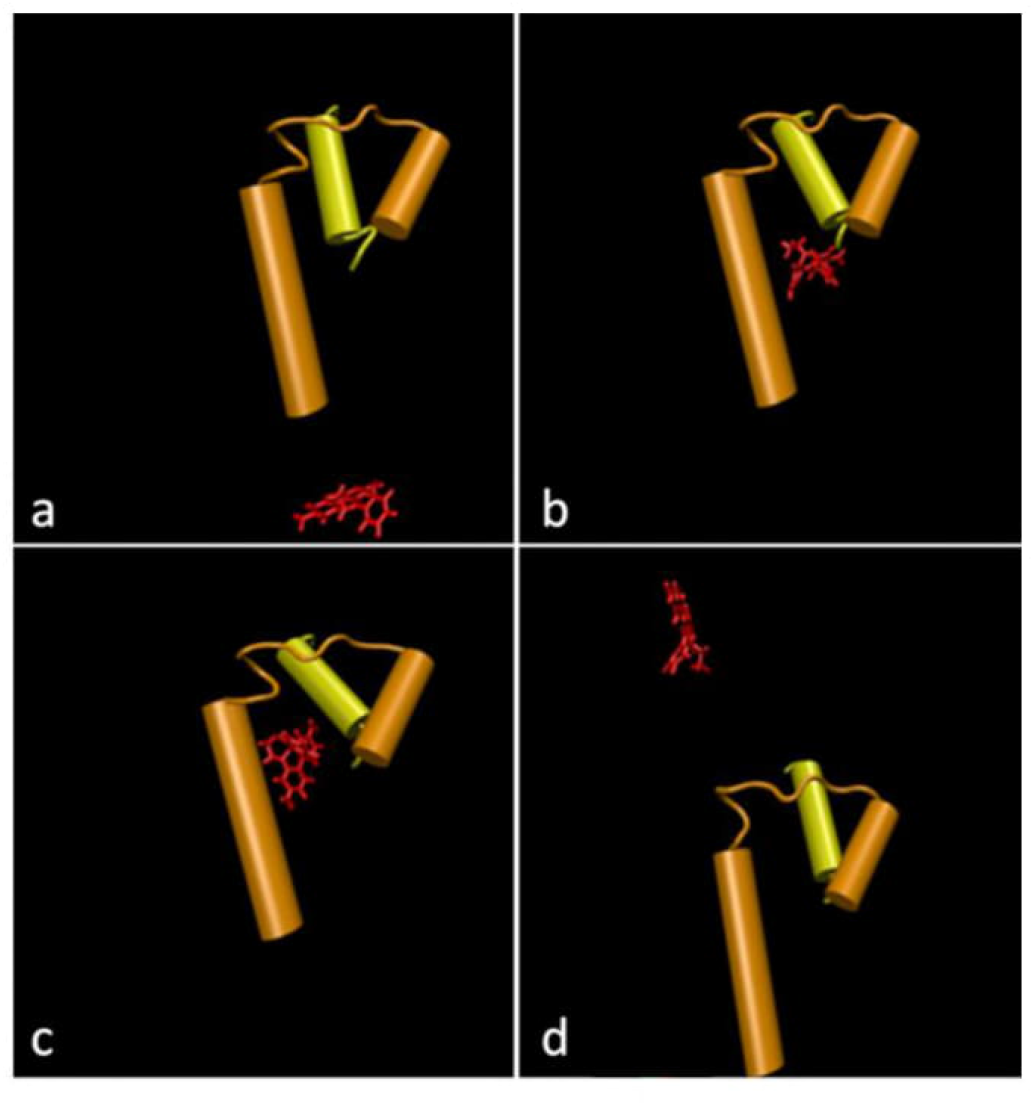
Detailed view of Helix 6 (yellow) shift during EtBr transit through Zone 1. Terminal sections of helix 1 and 2 (orange) are also shown, revealing their modest shifts during substrate transit. Panels a to d correspond to the corresponding images displayed in Fig. 5, i.e., panel a corresponds to position A, panels b and c are intermediate stages as EtBr moves through Zone 1, and panel d corresponds to position C.

Zone 2 (Fig. 3 and 7 in green) is composed primarily of β-strands and flexible loops, and is localized within the AcrB pore domain (10). The β-strand structure is quite stable and exhibits minimal structural fluctuation or positional displacement during the MD simulations, with the exception of one β-strand indicated by an arrow in Figs. 7 and 8a. The flexible loop and mobile β-strand residues are listed in Table 3. As the domain designation indicates, the β-strands form a well-defined pore connecting the transmembrane domain and the TolC interaction domain. Flexible loops 1 and 2 (Figs. 7 and 8) form a barrier or gate separating the transmembrane and pore domains during the MD simulations. However, when a ligand is present at position C, these loops undergo a conformational change that allows ligand passage into the pore domain. The loop conformational changes are accompanied by a shift of the amino-terminal end of one β-strand segment to permit ligand entry to the pore (Figs. 7 and 8). As the ligand traverses the pore region and reaches position D at the boundary of the pore domain and the TolC interaction domain, loops 1 and 2 assume their original “closed” conformation observed prior to ligand entry.

**Fig. 7.**
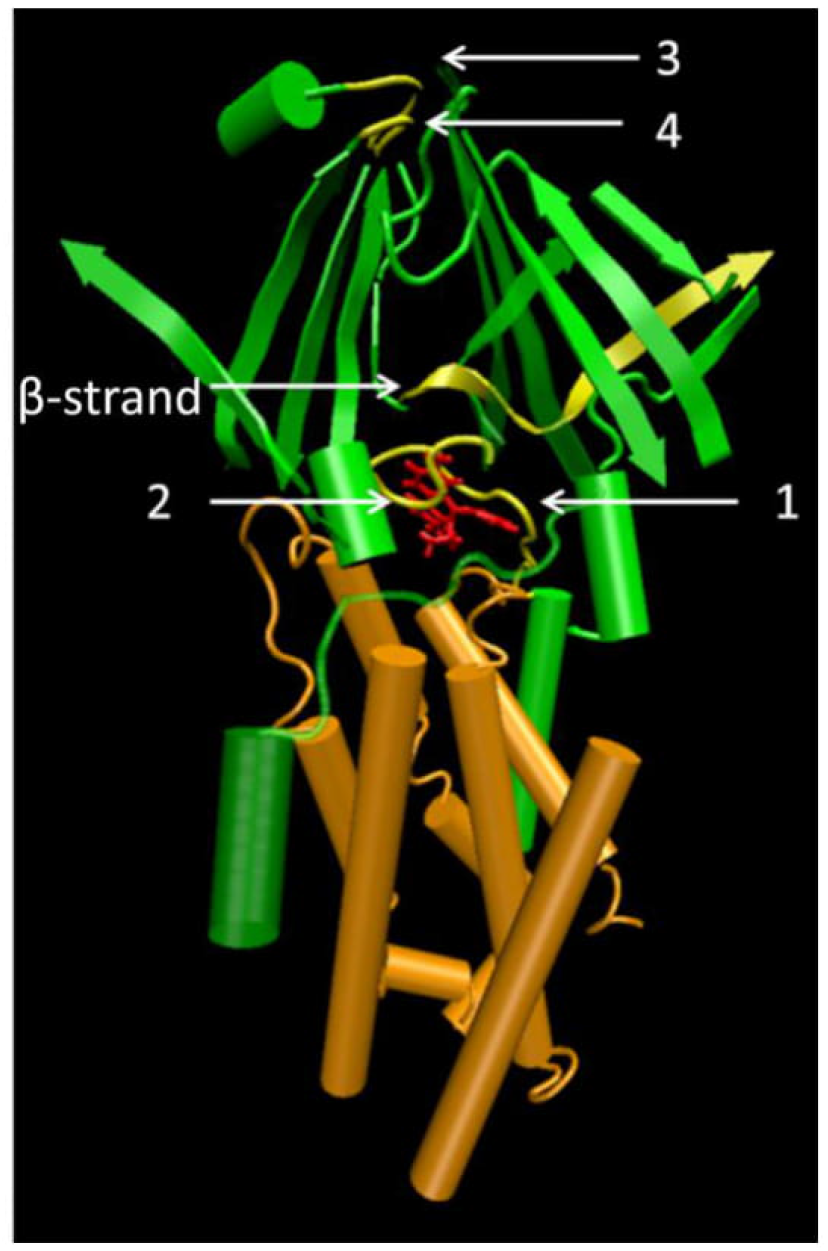
Zone 2, displayed in green, consists primarily of β-strands that form a well-defined pore. Loops 1 and 2 form a barrier or gate separating the pore domain (Zone 2) from the transmembrane domain (Zone 1). Flexible loops 3 and 4 form a barrier between the pore domain (Zone 2) and the TolC interaction domain (Zone 3; not shown in this Figure).

**Fig. 8.**
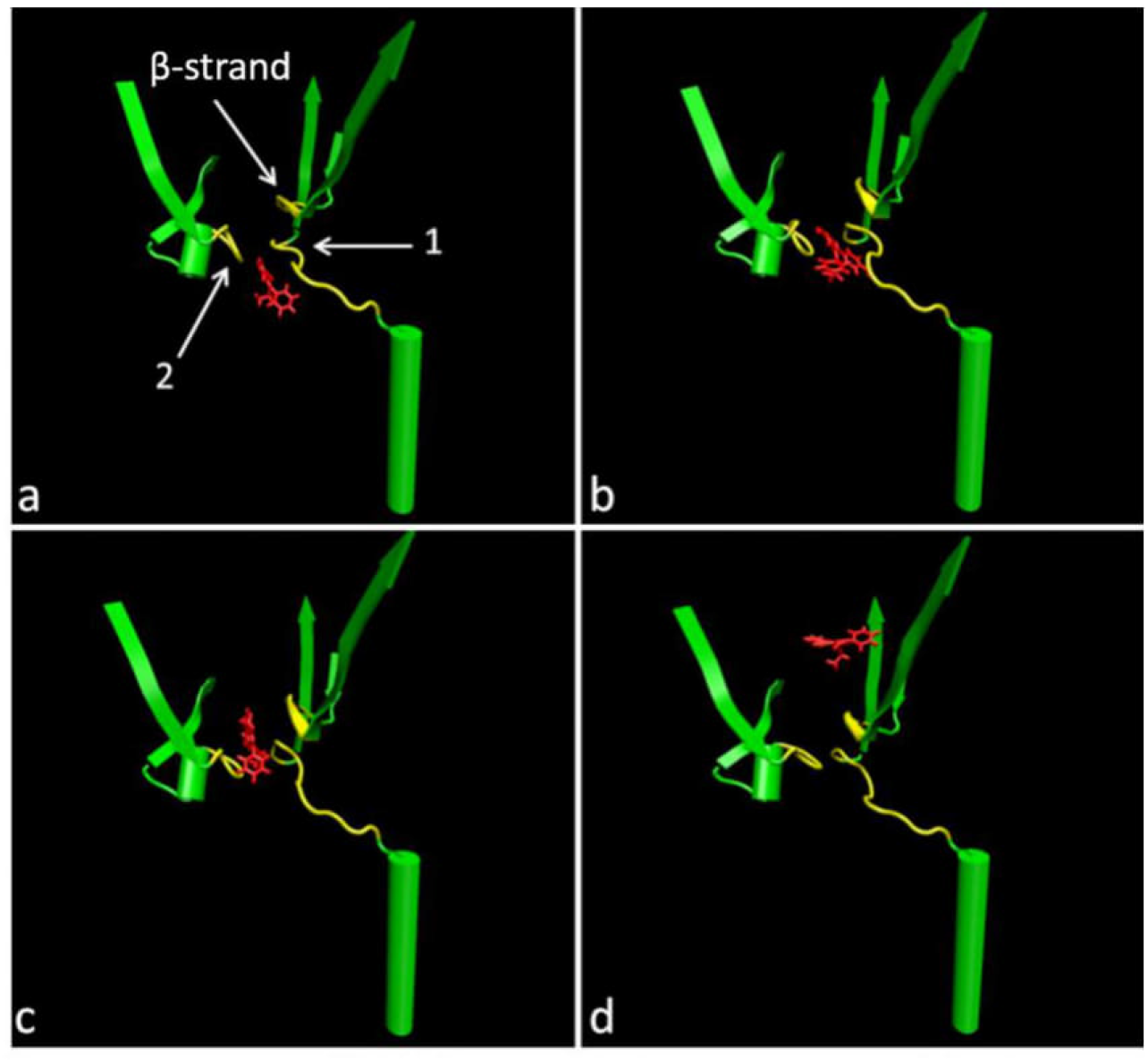
Detailed view of loop conformational changes and β-strand displacement observed as ligand traverses the pore from entry position C (panel 8a) to the exit point at position D (panel 8d).

**Table 3.** Residues of the loops and the β-strand that seemed to be of importance for the transport in Zone 2.

| Loop 1 | Loop 2 | β-strand | Loop 3 | Loop 4 |
| --- | --- | --- | --- | --- |
| 30 - 42 | 668 - 678 | 132 - 144 | 49 - 53 | 84 - 85 |

As the ligand approaches the boundary of the pore and TolC interaction domains (Fig. 7), loop 3 undergoes a significant conformational change and loop 4, immediately below loop 3, displays a modest conformational change (Fig. 9, panel a and b). These conformational changes allow the ligand to move into the TolC domain (Zone 3).

Zone 3, displayed in blue in Figures 3 and 9, consists of short helices and β-strands with numerous connecting loops in the TolC interaction domain. As the ligand moves from the pore domain into the TolC domain, helices 10 and 11 reorient to create an open passageway through the TolC domain. Loop 5 at the surface is quite flexible and undergoes a conformational change that allows the ligand to “escape” the AcrB protein into solution (i.e., the periplasmic space). While pathways for both ligands are nearly identical through Zones 1 and 2, the detailed features for EtBr and NUNL02 exit paths differ in Zone 3. NUNL02 rapidly exits Zone 3 to solution, while EtBr traverses the TolC domain interior and exits only after loop 5 opens sufficiently to allow passage (Fig. 9, Table 4).

**Fig. 9.**
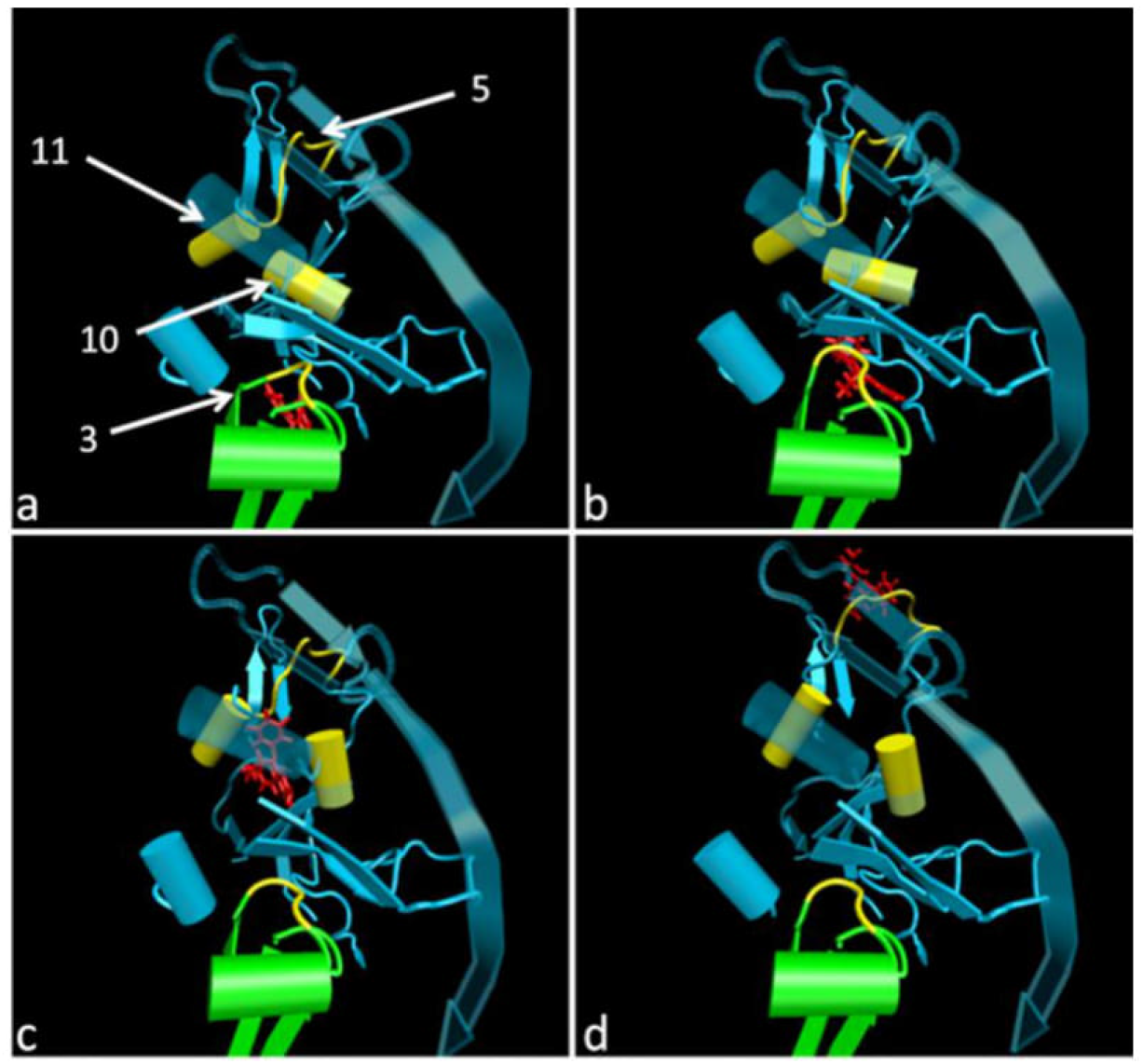
Zone 3. Loop 3 is at the border of Zone 2 and Zone 3. Helices 10 and 11 (yellow) display prominent displacements as the ligand moves through Zone 3. Loop 5 is an extremely flexible surface loop and presumably forms contacts with the TolC protein in the periplasm.

**Table 4.** Helix 10, 11 and loop 5 residues in Zone 3.

| Helix 10 | Helix 11 | Loop 5 |
| --- | --- | --- |
| 752-754 | 203-208 | 189-202 |

There is evidence to suggest that the TolC protein forms a complex with AcrB (24) (25). Loop 5 likely forms a portion of the AcrB-TolC interface, and explicit inclusion of the TolC protein in these models would clearly influence, and probably alter, details of the observed ligand transit pathways through the Zone 3 region.

#### 3.2.2 Pathway 2

The ligand entry point for pathway 2 is from the bilayer, as proposed previously by Nikaido *et al.* (20), unlike pathway 1 where ligands are captured directly from the cytoplasm. Position B identified in the docking calculations is used as the initial state for the pathway 2 TMD simulations (Fig. 10). The simulations reveal that ligands do not enter the transmembrane domain (Zone 1) directly from position B, but instead slide along the exterior of the transmembrane domain and penetrate the AcrB protein near position C (Fig. 10 and movies in A.12 and A.13).

**Fig. 10.**
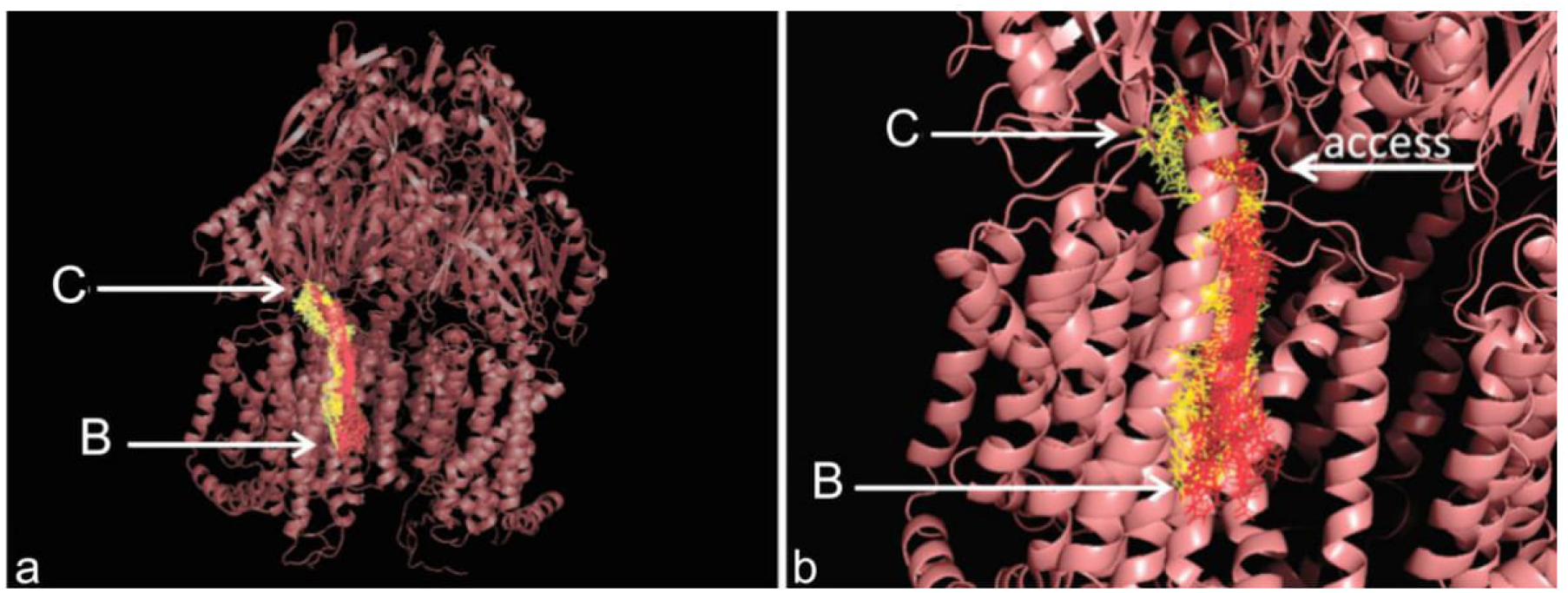
Zone 1 paths for ligand entry from the bilayer region at position B. a) The EtBr path is displayed in red and NUNL02 path in yellow. B) Detailed view of ligand paths shown in panel a.

There are no significant conformational changes or helix reorientations observed as ligands slide along the transmembrane domain helix bundle exterior. As the ligands approach the top of the helix bundle, loop 1 (Fig. 11) assumes an alternate conformation to allow ligand access to position C (the final state in this TMD segment). From position C, the remainder of the ligand efflux pathway is indistinguishable from pathway 1 described above.

**Fig. 11.**
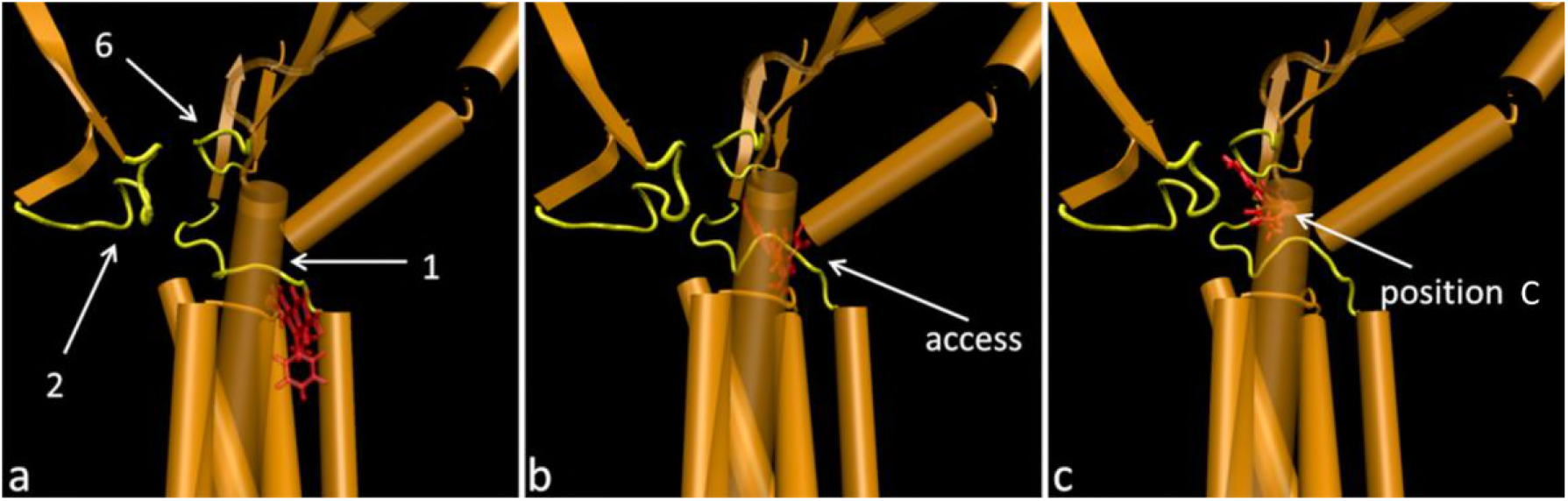
a) Loop structure at the interface of the transmembrane and pore domains. b) At the ligand entry point for pathway 2, loop 1 shifts to allow substrate entrance c) Loop 1 remains opened after the substrate reaches position C.

## 4. Discussion

The asymmetric configuration of the AcrB structure 4DX5 suggests an intriguing model for drug transport, based on conformational cycling of the monomers between loose, tight and open configurations (24) (11). However, in this study, we used the symmetric structure 1IWG and the TMD results suggest that each monomer may be able to function independently, capturing and conducting the substrate to the TolC protein for final extrusion. This independent monomer mechanism is much simpler as it does not require extensive interaction, e.g., “communication”, between the three monomers during the efflux process. This mechanism also implies greater efflux efficiency if all three monomers could function independently. However, our current calculations do not suggest in any way that the trimer cycling mechanism is not also plausible, and the actual efflux mechanism might involve components of both models.

In this work, our division of the AcrB protein into three distinct zones is based primarily on our molecular docking studies, which identified stable, intermediate biding sites for EtBr and NUNL02. However, we note that our domain or zone definitions correspond closely to the original structural characterization of three distinct domains (10). It seems unlikely that this close correspondence between structural and “functional” domain characterization is coincidence.

Independent proposals suggest that AcrB captures ligands directly from the cytoplasm (26) or from the outer leaflet of the cytoplasmic membrane (20). We explored both options in our current studies. Docking position A that functions as the starting point for efflux pathway 1 is consistent with direct ligand capture from the cytoplasm. This direct capture mechanism is simple and does not depend upon ancillary proteins, e.g., EmrR or MdfA, to transport ligands to the periplasmic space prior to AcrB capture (27).

The transmembrane domain is an α-helical bundle that forms a tunnel-like structure in the Zone 1 region of efflux pathway 1. The TMD results suggest that ligands can traverse this apparent tunnel passage with negligible energy barriers and only modest protein conformational changes. After the ligand has moved through Zone 1, the helical bundle quickly relaxes back to the starting protein conformation. We observed a peristaltic motion of the helical bundle as the ligand transits, but this motion is likely due to induced conformational changes caused by the ligand transit process. At present, we have no evidence that this peristaltic motion of the helical bundle is an intrinsic feature of the AcrB protein.

Docking position C is located in Zone 2, the AcrB pore domain, and is the most favorable ligand binding site in the entire protein identified in our docking calculations (Table 1). Access to position C (Fig. 10b) is controlled by conformational changes in loops 1 and 2 (Fig. 8) for efflux pathway 1, and conformational changes in loops 1, 2 and 6 (Fig. 11) for efflux pathway 2. Position C appears to correspond closely to a deep binding pocket described previously by Eicher *et al.* (17), and loops 1, 2 and/or 6 would correspond to the “switch-loops” they described that control access to the ligand binding pocket.

There is also a pair of flexible loops 3 and 4 (Fig. 12) that control the exit of ligand from Zone 2, the pore domain, to Zone 3, the TolC domain. Thus, there appears to be a clear “gating” mechanism for ligand entrance and exit in the pore domain. As noted above, once the ligand reaches Zone 3, exit from the AcrB protein is facile and rapid. Explicit inclusion of the TolC protein in the complex would certainly alter this final exit process, but we cannot speculate on the details based on our current calculations.

**Fig. 12.**
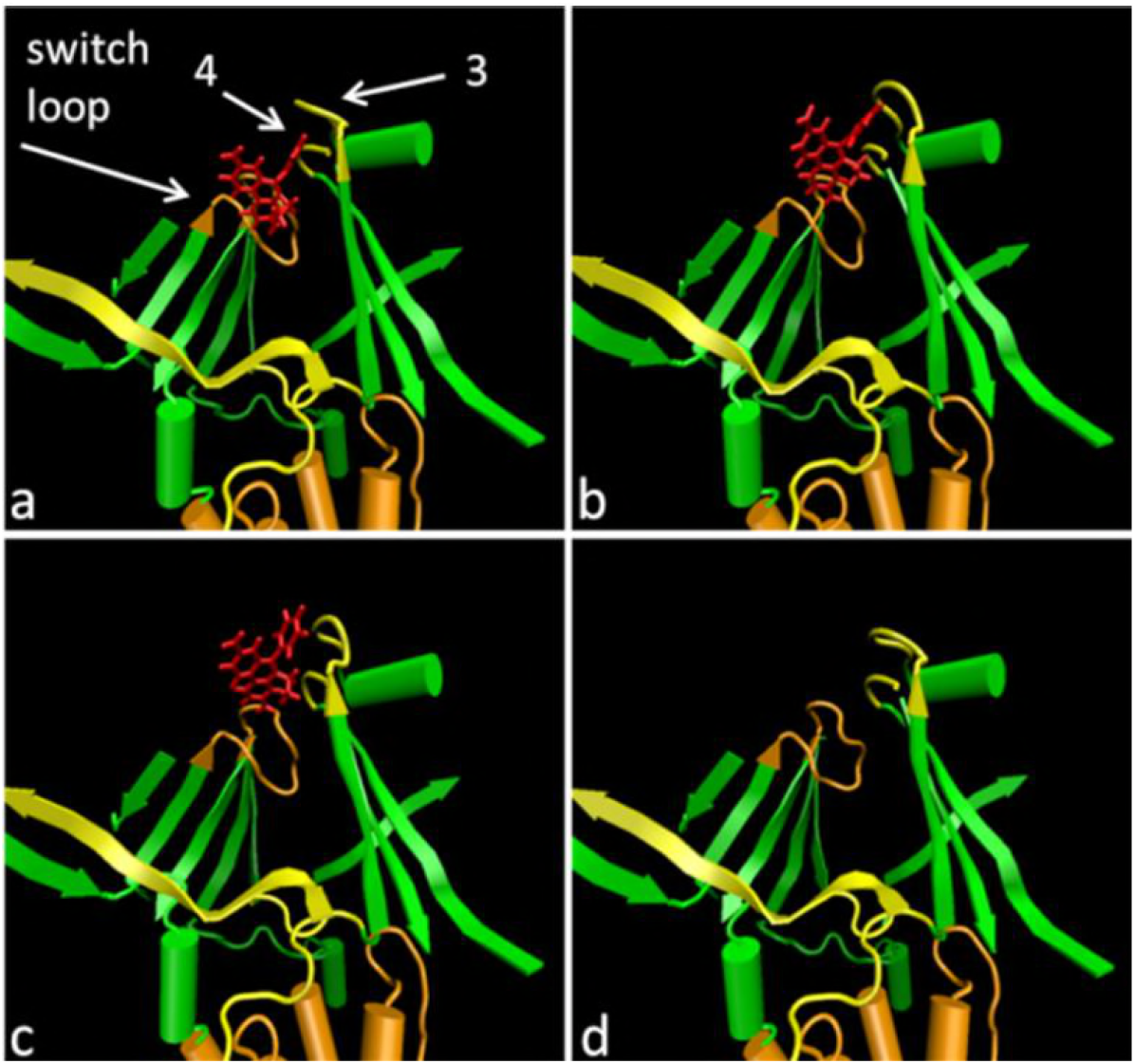
the switch-loop in orange [18] and loops 3 and 4. Notice how loop 3 lifts to allow the passage of the substrate, EtBr (licorice, in red). Loop 4 do not show a major displacement as loop 3, but it does move, as can be noticed comparing panels a & b.

Pathway 2 is interesting because position B, on the exterior surface of the transmembrane domain in the cytoplamsmic membrane outer leaflet region, is a plausible capture point for nonpolar ligands that might localize in the membrane. TMD results suggest that the substrate slides along the helical bundle surface until it finds an entrance point and reaches position C. Loops 1, 2 and 6 (Fig. 11) control access for ligands from the helix bundle exterior in the outer leaflet region to position C in the pore domain. The possibility of lateral capture of substrates from the outer leaflet of the cytoplasmic membrane is intriguing and might have a favorable impact on efflux efficiency. The substrate would have a larger area to dock, rather than a small, specific binding site, e.g., position A in Zone 1. Of course, it is possible that some ligands might prefer pathway 1 and others pathway 2, e.g., as a function of ligand lipophilicity, etc.

The TMD results suggest that, for both pathways, the substrates EtBr and NUNL02 follow very similar efflux trajectories until position D, where the two ligand display dramatically different exit trajectories (Fig. 2). As noted above, explicit inclusion of the TolC protein in the complex would undoubtedly alter the exit pathway details from position E substantially for both ligands. It is known that NUNL02 has high affinity for AcrB, and these simulation results support the possibility of efflux inhibition by competition between substrates as proposed previously (9). Thus, successful efflux inhibitor design may not require development of molecules that block drug binding at key sites or entry points via direct competitive binding, but simply discovery of molecules that follow similar efflux trajectory pathways, thus diminishing drug efflux by saturating the transport path, effectively creating a “traffic jam”.

Finally, the TMD results showed no evidence that substrates might be extruded through the AcrB central pore, in good agreement with a previous study (26).

## 5. Conclusion

The technique of Targeted Molecular Dynamics was used to study how EtBr and NUNL02 might be captured and transported by the AcrB efflux pump. The simulations were performed for two distinct efflux pathways, based on two difference proposals for substrate capture, and revealed that loops 1, 2 and 6 (border of Zones 1 and 2), 3 and 4 (border of Zones 2 and 3), and 5 (Zone 3), α-helices 1 to 9 (Zone 1), 10 and 11 (Zone 3) and a β-strand (residues 132-144, Zone 2) play an active role in transport, interacting extensively with the substrates as they were extruded. Further, the simulations suggested that EtBr and NUNL02 would compete for the same efflux routes, regardless of specific pathway. This finding suggests that the mechanism of efflux inhibition by competition between molecules is plausible. These calculations provide molecular details for plausible substrate efflux pathways and suggest a number of new biophysical experiments to address the AcrB efflux mechanism in greater detail.

## Acknowledgements

Lande Silva Jr wishes to thank Dr. Jonathan Sheeham, for his teachings in molecular dynamics, and all the staff of the Center for Structural Biology from Vanderbilt University, TN, USA.

## Funding

This work was supported by the Coordenação de Aperfeiçoamento de Pessoal de Nível Superior/Ministério da Educação (CAPES/MEC), Brazil, doctoral grant to Lande Silva Jr and Karina S. Machado acknowledges CAPES “edital biologia computacional número 051/2013”.

Conflict of interests: None declared.

Ethical approval: Not required.

## Supplementary Figures

**Fig. A1:**
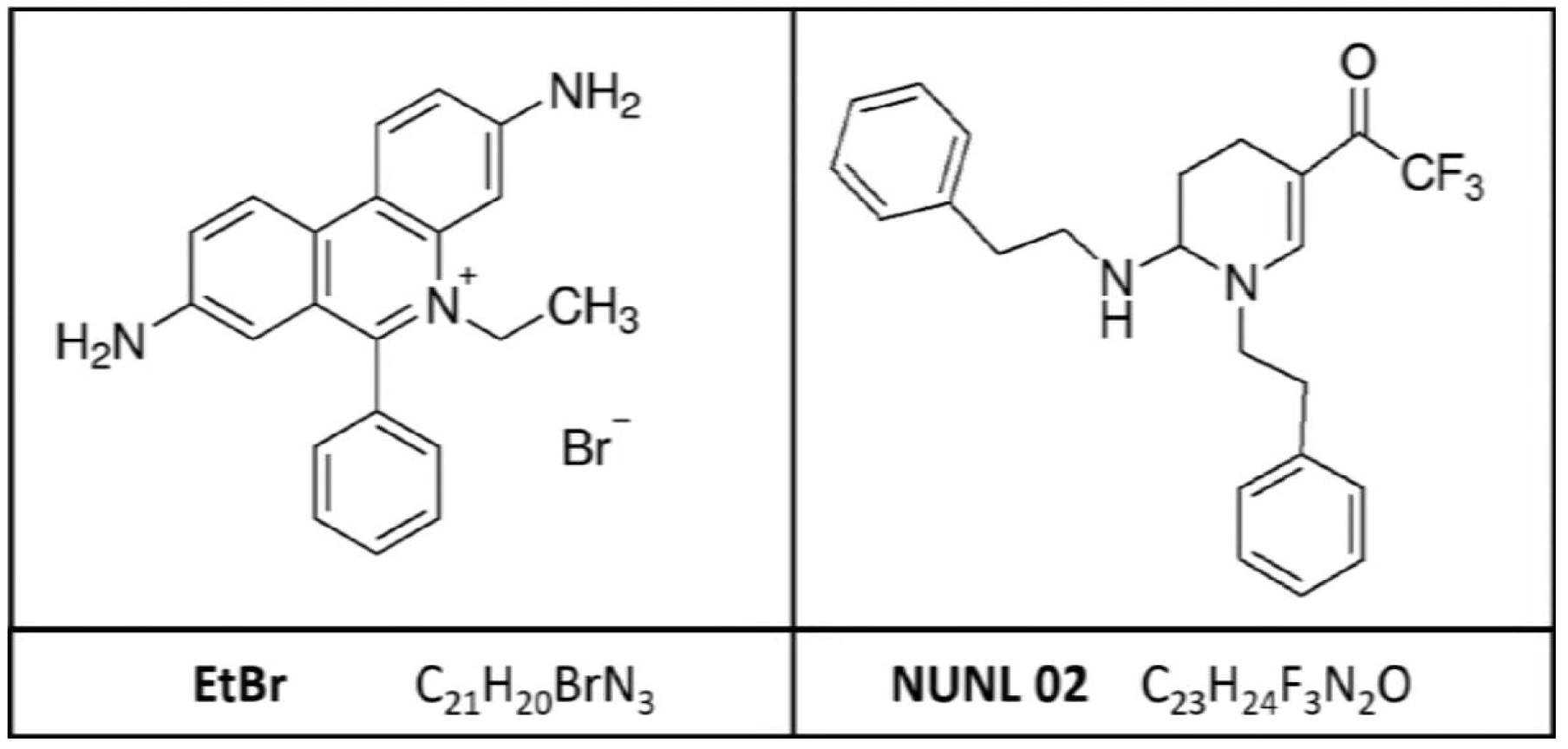
structural and molecular formula of the ethidium bromide (left, upper and lower panel) and a tetrahydropyridine derivative, NUNL02, (right, upper and lower panel), respectively.

**Fig. A2:**
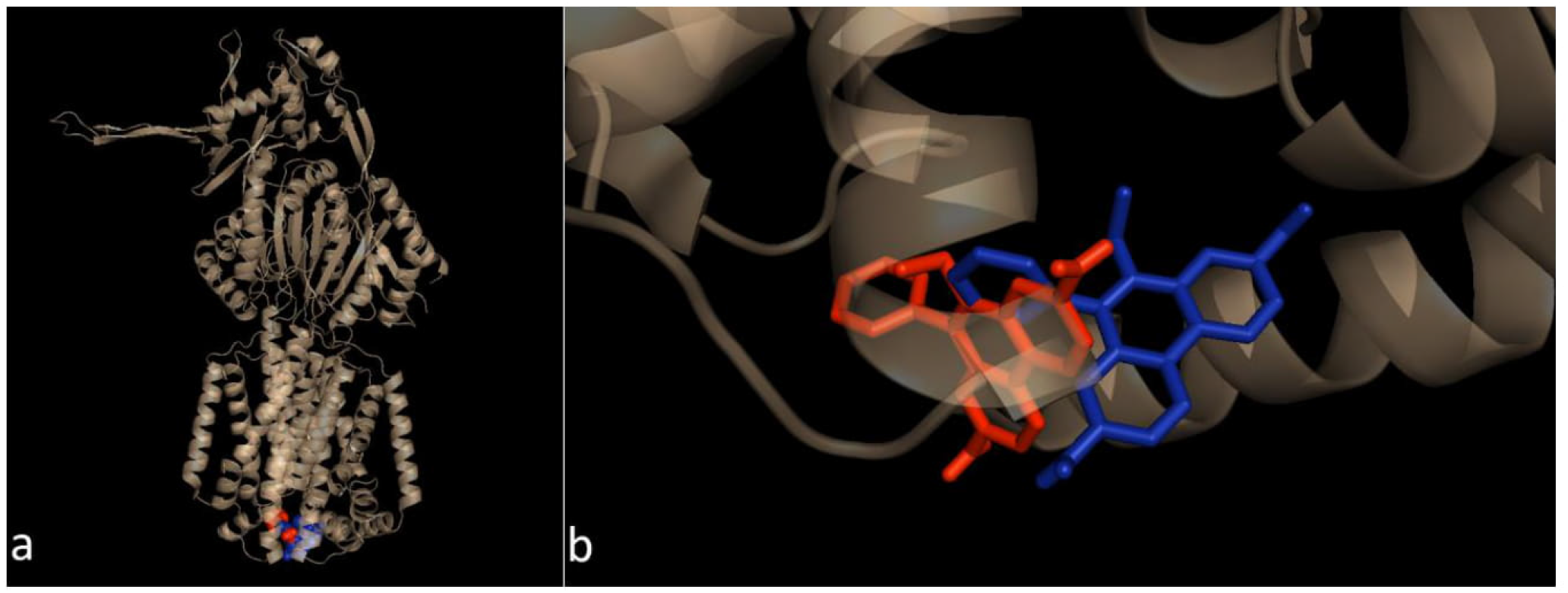
Docking equivalences. a) EtBr in red spheres is in position A (FEB of -5.8 kcal/mol (exhaustiveness 8, grid 40 × 48 × 7). EtBr in blue spheres is in the alternate place found (second best pose, the first one was of a FEB of -6.7 kcal/mol and did not have superposition), when we increased the size of the grid and the exhaustiveness (FEB of -6.4 kcal/mol, exhaustiveness 128, grid 65 × 65 × 23). b) There is just a slight difference in the spatial orientation between the two dockings, but the proximity of the positions is evident.

**Fig. A3:**
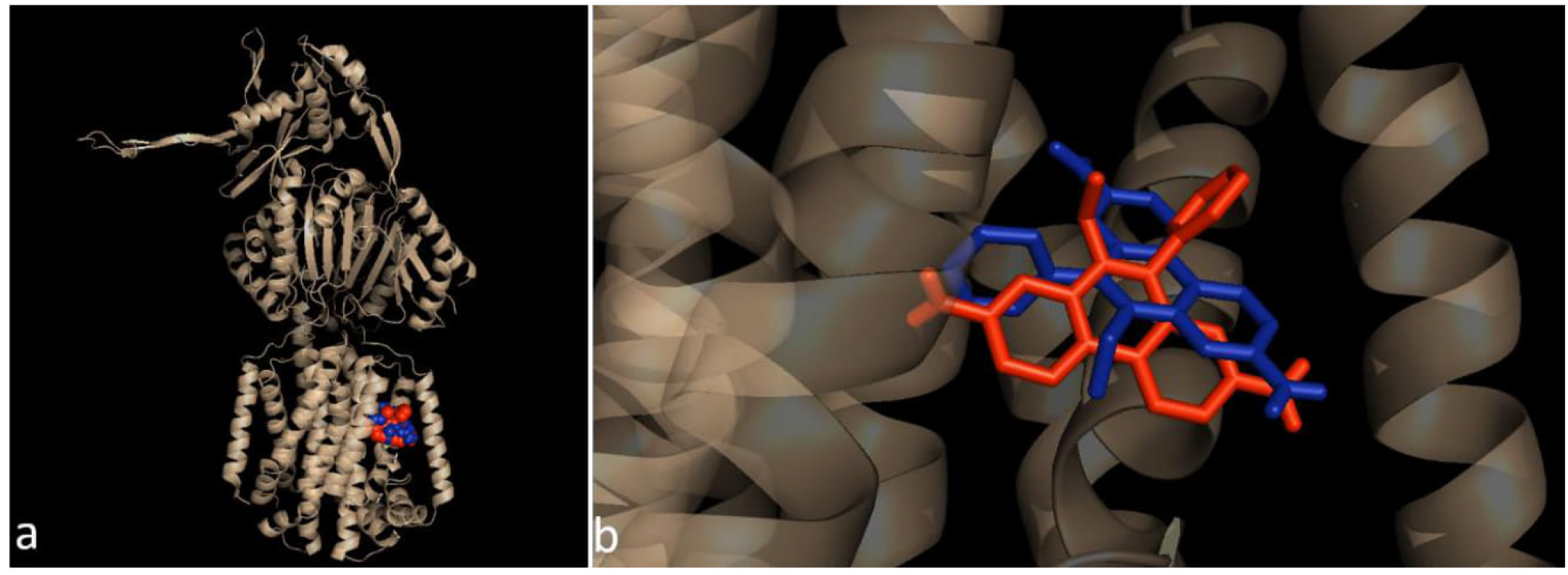
Docking equivalences. a) EtBr in red spheres is in position B (FEB of -7.1 kcal/mol (exhaustiveness 8, grid 40 × 48 × 7). EtBr in blue spheres is in the alternate place found when we increased the size of the grid and the exhaustiveness (FEB of -7.6 kcal/mol, exhaustiveness 128, grid 65 × 65 × 23). b) The EtBr in red (position B) and blue (alternate) sticks, for the docking variations, are occupying the same site, in a slightly diferent orientation.

**Fig. A4:**
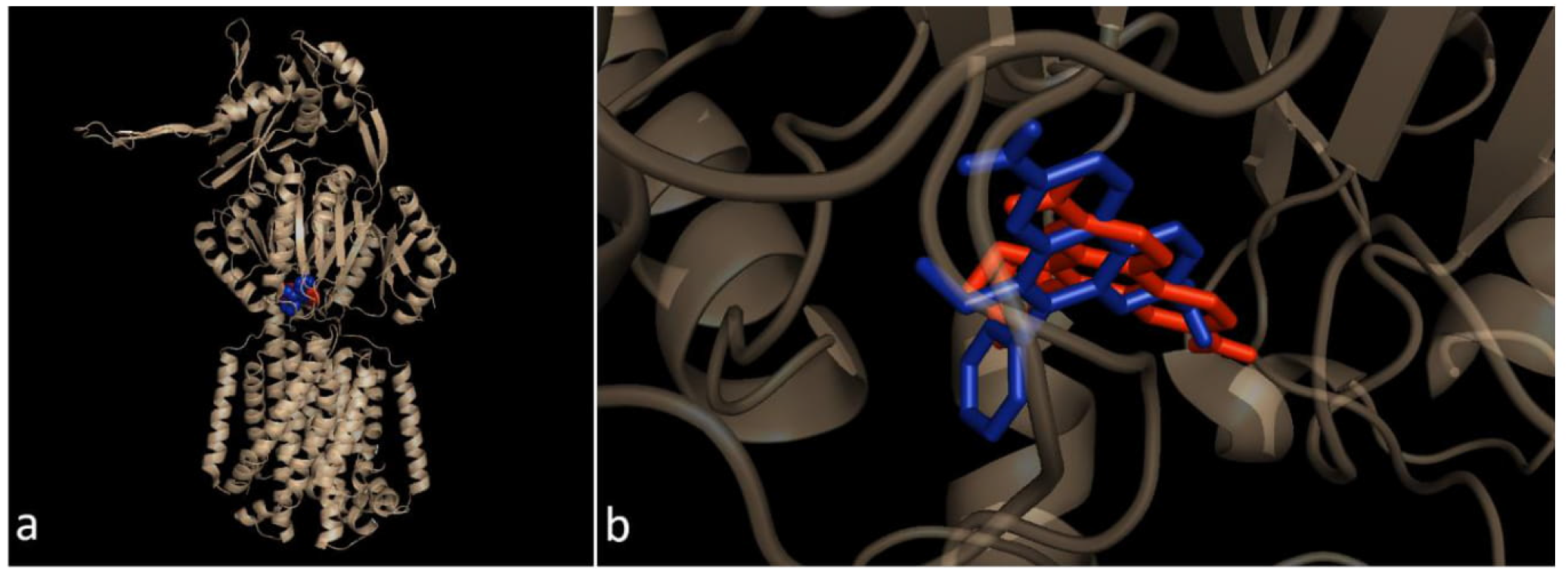
Docking equivalences. a) EtBr in red spheres is in position B (FEB of -8.2 kcal/mol (exhaustiveness 8, grid 40 × 48 × 7). EtBr in blue spheres is in the alternate place found when we increased the size of the grid and the exhaustiveness (FEB of -8.7 kcal/mol, exhaustiveness 128, grid 65 × 65 × 23). b) The EtBr in red (position B) and blue (alternate) sticks, for the docking variations, are occupying the same site, in a slightly diferent orientation.

**Fig. A5:**
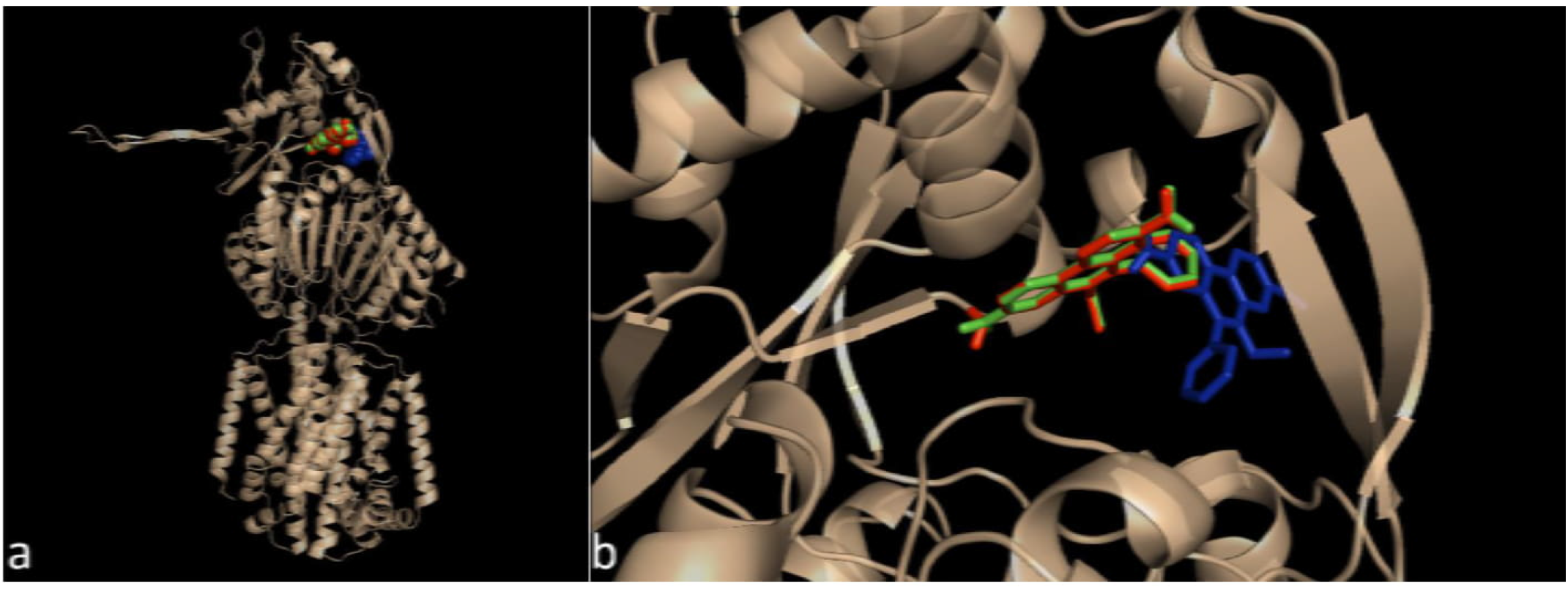
Docking equivalences. a) EtBr in red spheres is in position D (FEB of -7.4 kcal/mol (exhaustiveness 8, grid 40 × 48 × 7). EtBr in blue spheres is in the alternate place found for pose 1, when we increased the size of the grid and the exhaustiveness (FEB of -7.6 kcal/mol, exhaustiveness 128, grid 65 × 65 × 23), finally, the EtBr in green is the second best pose for the alternate position. b) The EtBr in red (position B) and blue (alternate) sticks, for the docking variations, are occupying the same pocket, with some superposition, however, the second best pose in green sticks (FEB of -7.4 kcal/mol) coincides with position D.

**Fig. A6:**
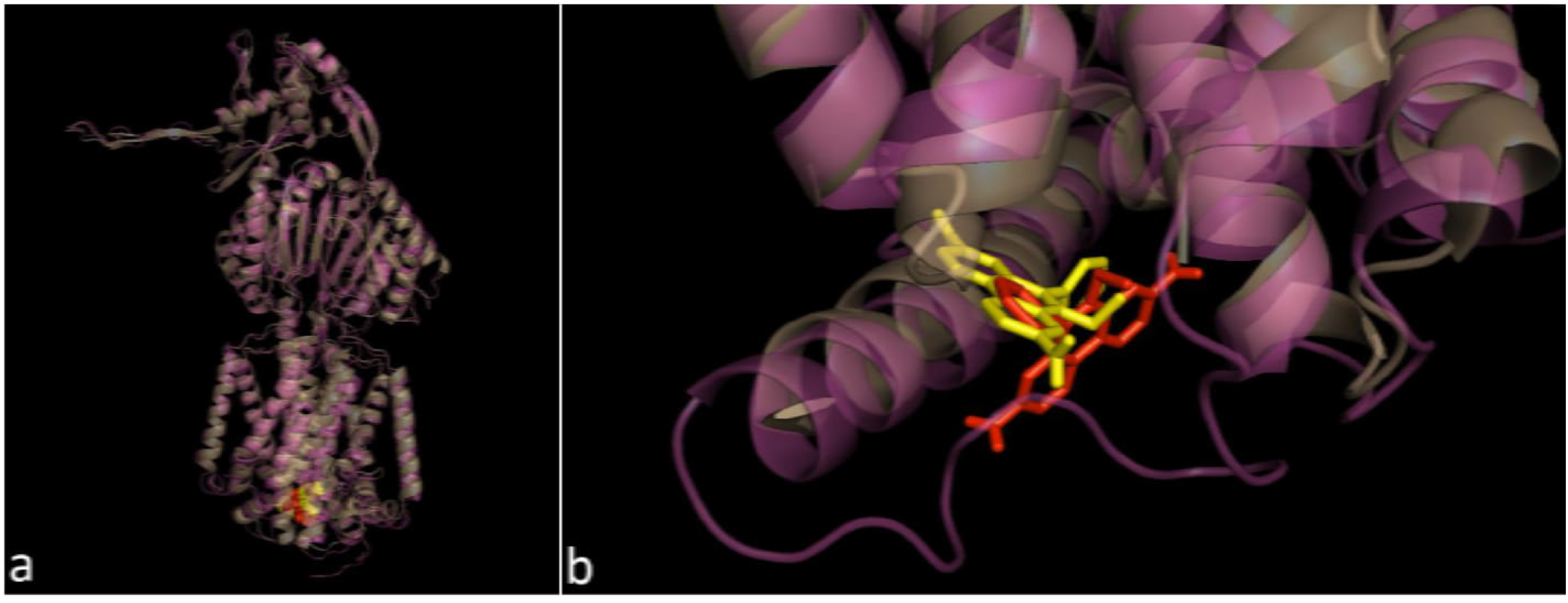
Docking equivalences for EtBr in structures 1IWG (gray cartoon) and 4DX5-A (magenta). a) EtBr in red spheres is in position A (FEB of -5.8 kcal/mol (exhaustiveness 8, grid 40 × 48 × 7). EtBr in yellow spheres is in the best docking position in structure 4DX5-A (FEB of -9.4 kcal/mol, exhaustiveness 128, grid 65 × 65 × 23). b) EtBr in red sticks, for position A and in yellow sticks for the docking in structure 4DX5-A, notice the superposition between them.

**Fig. A7:**
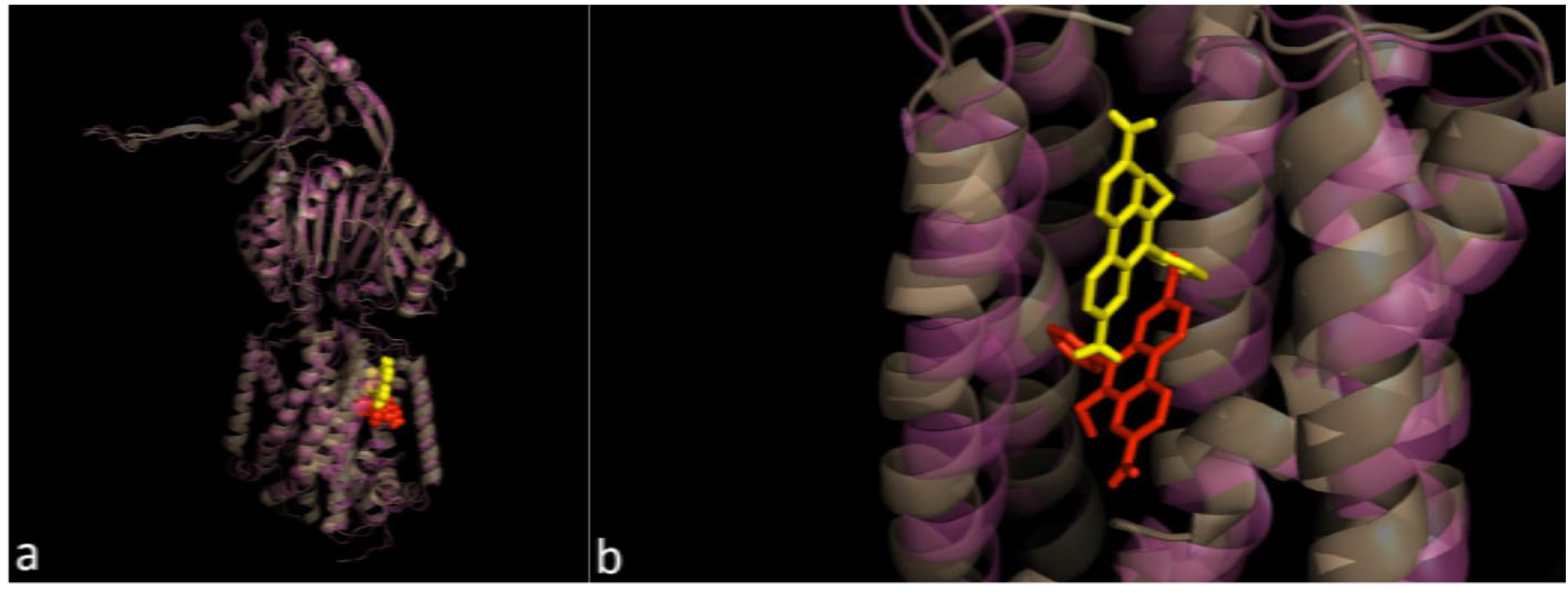
Docking equivalences for EtBr in structures 1IWG (gray cartoon) and 4DX5-A (magenta). a) EtBr in red spheres is in position B (FEB of -7.1 kcal/mol (exhaustiveness 8, grid 40 × 48 × 7). EtBr in yellow spheres is in the best docking position in structure 4DX5-A (FEB of -7.8 kcal/mol, exhaustiveness 128, grid 65 × 65 × 23). b) EtBr in red sticks, for position B and in yellow sticks for the docking in structure 4DX5-A, notice the EtBr for both dockings is between the same beta strands, with some superposition, although EtBr in 4DX5-A is a little above in the figure.

**Fig. A8:**
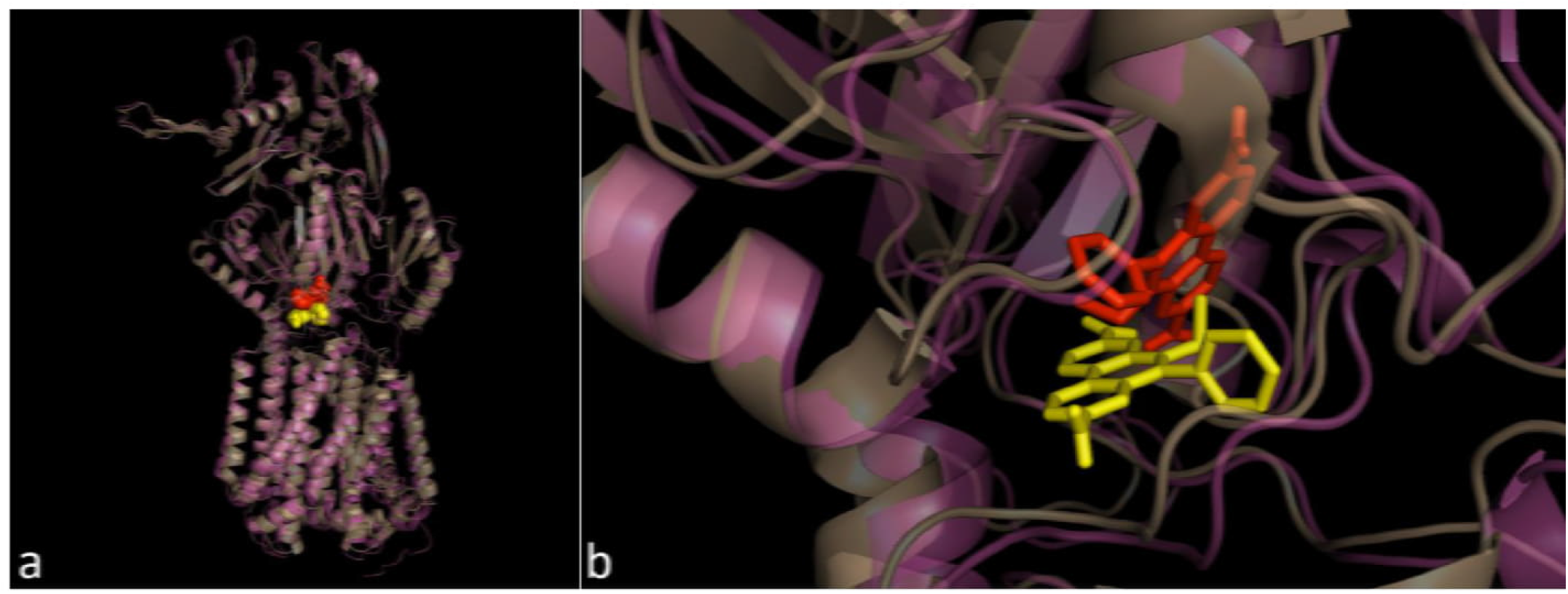
Docking equivalences for EtBr in structures 1IWG (gray cartoon) and 4DX5-A (magenta). a) EtBr in red spheres is in position C (FEB of -8.2 kcal/mol (exhaustiveness 8, grid 40 × 48 × 7). EtBr in yellow spheres is in the best docking position in structure 4DX5-A (FEB of -8.1 kcal/mol, exhaustiveness 128, grid 65 × 65 × 23). b) EtBr in red sticks, for position C and in yellow sticks for the docking in structure 4DX5-A, the EtBr representations are between the same loops for both dockings, very close with a diferent spatial orientation and some superposition.

**Fig. A9:**
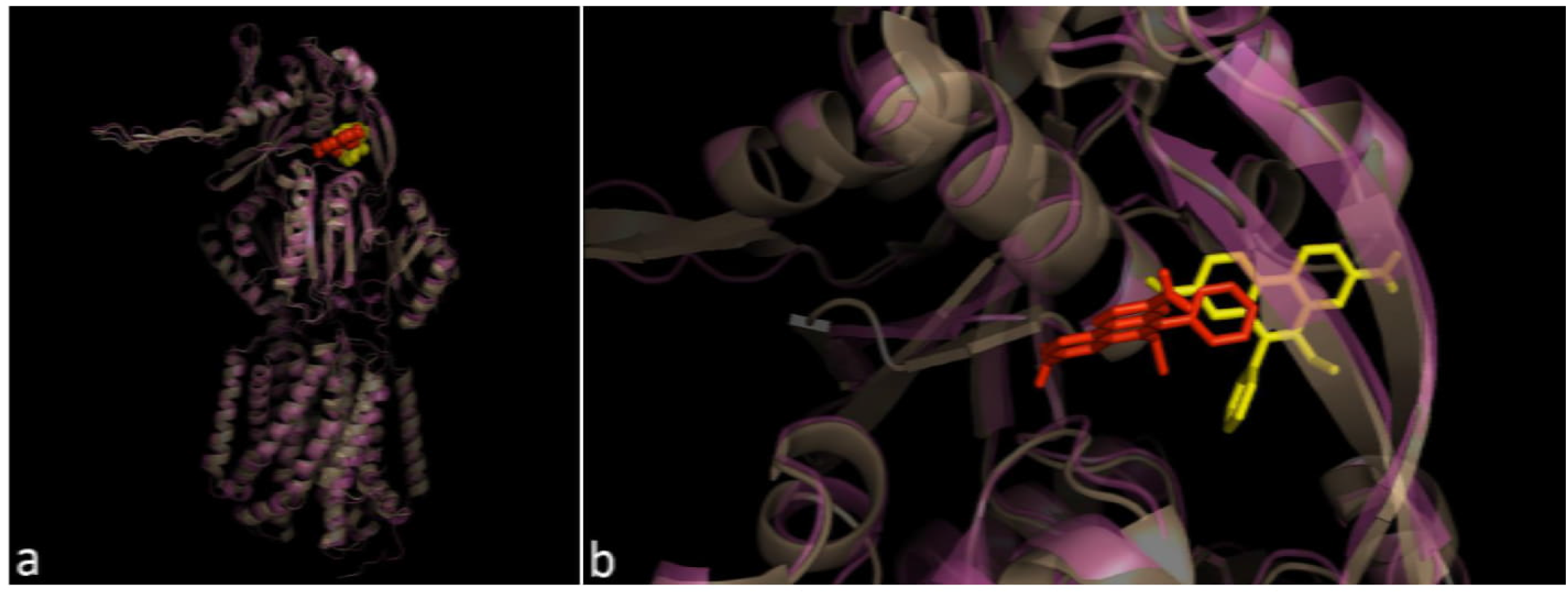
Docking equivalences for EtBr in structures 1IWG (gray cartoon) and 4DX5-A (magenta). a) EtBr in red spheres is in position C (FEB of -7.4 kcal/mol (exhaustiveness 8, grid 40 × 48 × 7). EtBr in yellow spheres is in the best docking position in structure 4DX5-A (FEB of -7.6 kcal/mol, exhaustiveness 128, grid 65 × 65 × 23). b) EtBr in red sticks, for position C and in yellow sticks for the docking in structure 4DX5-A, the EtBr representations are in the same pocket for both dockings, very close with a diferent spatial orientation and some superposition.

